# Neural dynamics of illusory tactile pulling sensations

**DOI:** 10.1101/2021.10.12.464029

**Authors:** Jack De Havas, Sho Ito, Sven Bestmann, Hiroaki Gomi

**Affiliations:** NTT Communication Science Laboratories, Japan; UCL Queen Square Institute of Neurology Department of Clinical and Movement Neurosciences, University College London, UK; Wellcome Centre for Human Neuroimaging, UCL Queen Square Institute of Neurology, University College London, UK

**Keywords:** asymmetric vibration, somatosensory, SEP, N140, P200, P3b, tactile illusion, parietal lobe, tangential force

## Abstract

The sensation of directional forces and their associated sensorimotor commands are inextricably intertwined, complicating the identification of brain circuits responsible for tactile pulling sensations. One hypothesis is that, like tactile frequency discrimination, pulling sensations are generated by early sensory-frontal activity. Alternatively, they may be generated later in the somatosensory association cortex. To dissociate these accounts and uncouple the pulling sensation from unrelated but correlated sensory and motor processing, we combined high-density EEG with an oddball paradigm and asymmetric vibration, which creates an illusory sensation of the hand being directionally pulled. Oddballs that created a pulling sensation in the opposite direction to common stimuli were compared to the same oddballs in the context of neutral common stimuli (symmetric vibration) and to neutral oddballs. Brain responses to having directional pulling expectations violated by directional stimuli were therefore isolated. Contrary to the sensory-frontal account, frontal N140 brain activity was actually larger for neutral than pulling oddballs. Instead, pulling sensations were associated with amplitude and latency modulations of midline P200 and P3b potentials, and specifically, to contralateral parietal lobe activity 280ms post-stimulus. The timing of this activity suggested pulling sensations involve spatial processing, such as tactile remapping between coordinate frames. Source localization showed this activity to be centered on the postcentral sulcus, superior parietal lobule and intraparietal sulcus, suggesting that pulling sensations arise via the processing of body position, tactile orientation and peripersonal space. Our results demonstrate how tactile illusions can uniquely disambiguate parietal contributions to somatosensation by removing unrelated sensory processing.

**Significance statement:** The neural mechanisms of tactile pulling sensations are poorly understood. Competing early sensory-frontal and later somatosensory association cortex accounts are hard to dissociate due to confounding sensory and motor signals present when forces are applied to the skin. Here, we used EEG and a novel asymmetric vibration approach to induce an illusory pulling sensation, which circumvents these issues. We found that pulling sensations were associated with parietal lobe activity 280ms post-stimulus and modulations of the P200. The timing and location of this activity suggested that pulling sensations necessitate spatial processing and supported a somatosensory association cortex account of the pulling sensation.

## Introduction

The sensation of directional force is vital in everyday life, allowing us to, for example, know our dance partner’s intention, or quickly learn the physical properties of a touched object (Johansson and Flanagan, 2009). Despite much progress in understanding peripheral tactile processing (Johansson et al., 1992a; Panarese and Edin, 2011; Pruszynski and Johansson, 2014; Pruszynski et al., 2018), little is known about how pulling sensations arise in the human brain. Research using monkeys shows that tangential forces are processed rapidly (~50ms post-stimulus) in the primary somatosensory cortex (SI) (Salimi et al., 1999; Fortier-Poisson and Smith, 2016; Fortier-Poisson et al., 2016). However, it is unclear if such processing is sufficient to give rise to directional pulling sensations.

Pulling sensations likely require activity beyond SI. A circuit involving SI, SII and the prefrontal cortex (PFC) has been demonstrated to underpin the perception of tactile frequency (Romo and Salinas, 2003; de Lafuente and Romo, 2006; Hernández et al., 2010). If the discrimination of directional pulling sensations requires only the accessing and comparing of stored patterns of activity in SI and SII, then these, or closely related, sensory-frontal circuits may be sufficient. If, however, directional pulling sensations necessitate spatial processing (Badde and Heed, 2016), then the parietal cortex may play an important role. On this account, activity in superior parietal lobule (SPL) and intraparietal sulcus (IPS) combine body position information with information about the spatial direction of the force to generate a directional pulling sensation (Ehrsson et al., 2003; Van Boven et al., 2005; Sack, 2009).

Determining precisely when pulling sensations emerge will constrain mechanistic accounts. Pulling sensations are assumed to depend on force vector extraction. If this extraction occurs during initial feedforward processing in SI and SII, early neural correlates, such as N140 enhancement, are expected. The N140 originates in SII and the PFC (Desmedt and Tomberg, 1989; Frot et al., 1999). It is a reliable marker for tactile awareness (Auksztulewicz et al., 2012; Schröder et al., 2021) and texture processing (Genna et al., 2018). Further, the N140 is modified by exogenous and endogenous attention (Nakajima and Imamura, 2000), so if pulling sensations emerge upstream, indirect, attention-related N140 enhancement should be observed. Conversely, if pulling sensations require later spatial processing, then the P200 or P3b instead will be enhanced. The pulling sensation may depend on mapping the force vector from skin centered to external coordinates, known as tactile remapping (Driver and Spence, 1998; Heed et al., 2015). Tactile remapping occurs after initial somatosensory processing and has been linked to the P200 (Longo et al., 2012; Bufalari et al., 2014).

Determining the neural mechanisms of pulling sensations has been difficult because traditional stimuli, such as active touch, sudden loads applied to held objects or tangential forces applied passively to the skin, are accompanied by correlated but unrelated motor and sensory processing (Johansson et al., 1992b; Birznieks et al., 2001). To disambiguate pure sensations of pulling from their conjoined sensory and motor processes, we here use an asymmetric vibration approach, which creates a strong, illusory sensation of being pulled in a particular direction via a small handheld device, without active movement (Amemiya et al., 2005; Amemiya and Maeda, 2008; Tappeiner et al., 2009; Amemiya and Gomi, 2014, 2016; Tanabe et al., 2018; Gomi et al., 2019). Symmetric vibration can be used as a control stimulus, which is closely matched in terms of stimulus complexity, but does not induce an illusory pulling sensation.

We recorded high-density EEG while participants performed a tactile oddball task in which uncommon target stimuli must be detected from a stream of common stimuli (Shinozaki et al., 1998; Kida et al., 2003; Spackman et al., 2007). Oddballs that created an illusory pulling sensation in the opposite direction to the common stimuli (asymmetric vibration) were compared to the same oddballs in the context of neutral common stimuli (symmetric vibration), and also to neutral oddball stimuli. These relative oddball effects meant we could isolate the brain activity specific to having directional expectations contradicted by directional stimuli, and therefore determine when and where the pulling sensation emerges in the brain, helping to dissociate spatial, parietal cortex accounts from non-spatial sensory-frontal accounts.

## Methods

### Equipment

Participants were seated at a table approximately 40cm from a computer monitor with their right forearm resting on an adjustable arm rest (Fig. 1D.). View of the right arm was obscured by a dividing screen. Symmetric and asymmetric vibration stimuli were delivered by a small, coin-sized device (Amemiya et al., 2005; Amemiya and Gomi, 2014) covered with grip tape (sandpaper grit density = #400) that was held between index finger and thumb in a pinch grip (Fig. 1B.). An accelerometer (356A03, PCB Piezotronics, Inc., New York, USA; sampling frequency = 4000Hz) was attached to the device. Accelerometer signals were displayed to the experimenter via an oscilloscope (TDS2004C, Tektronix, Inc., Oregon, USA), for the purposes of checking that the correct conditions were being administered at all times. Accelerometer signals were also recorded so that the precise stimulus onset time could be determined for every trial. Participants wore earplugs throughout the experiment to prevent auditory cues relating to the vibration conditions. Vibration onset timing, accelerometer recording, task instructions and fixation crosses were controlled via MATLAB (2017a) and Psychtoolbox (Brainard, 1997). Visual stimuli were displayed via a flat screen monitor (27-inch LCD, 1902 x 1080 pixels, 60 Hz refresh rate). EEG data were acquired via a 129 electrode net (HydroCel GES 300, MagstimEGI, Oregon, USA). Data were acquired at 1000Hz and Net Station EEG software (Magstim EGI, Oregon, USA).

**Figure 1:**
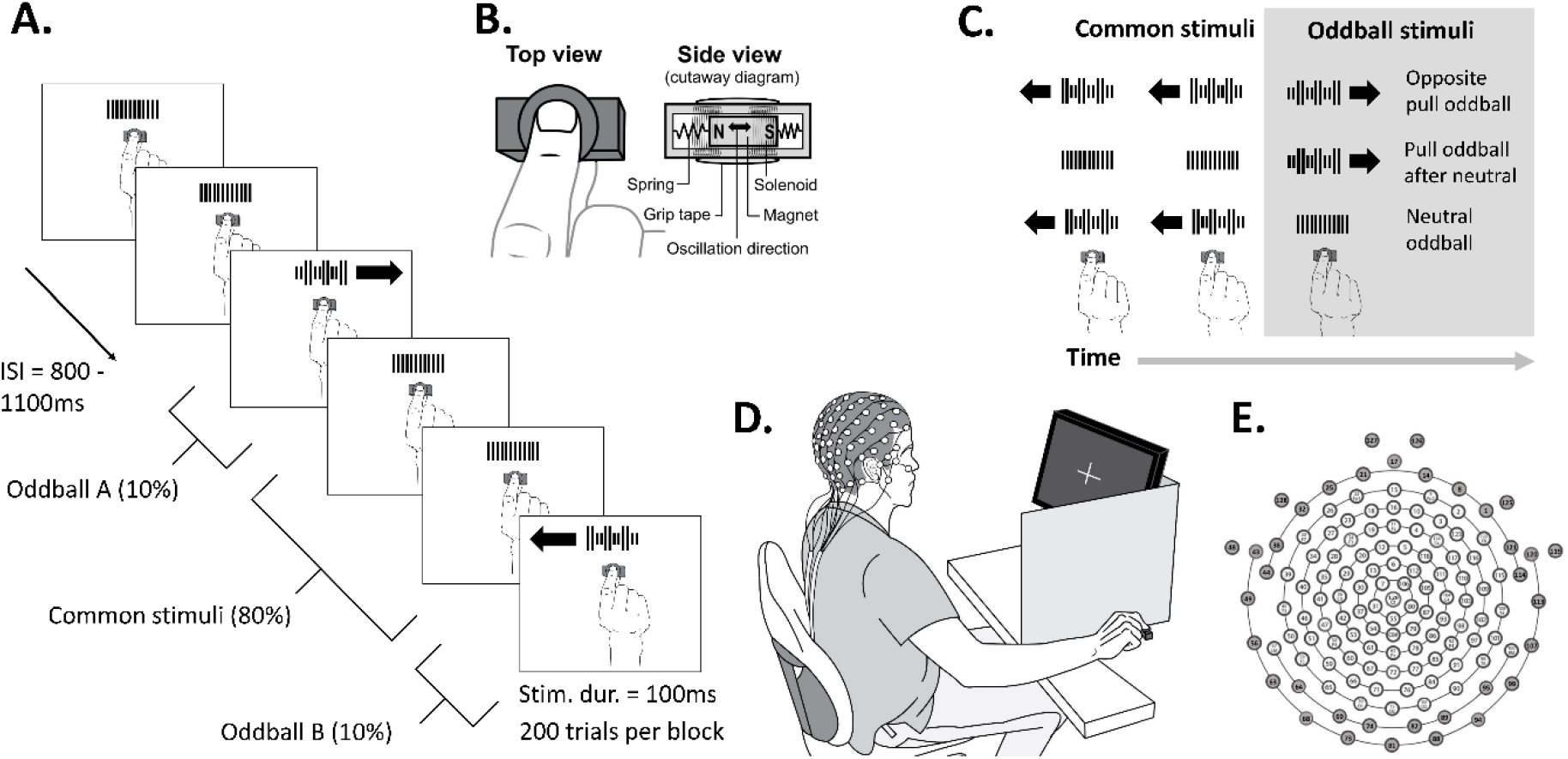
Task design and experimental setup. **A.** Task design showing a single block in which Neutral (symmetric vibration) is the common stimulus and Left and Right pull (asymmetric vibration) are the randomly appearing Oddballs. Note that there were two other types of blocks, in which the Left and Right pulling stimuli acted as the common stimuli, with the oddballs being Neutral and Right, and Neutral and Left respectively. Participants had to silently count all oddballs and report their count at the end of the block. **B**. Close up of the device used to generate asymmetric and symmetric vibration, showing top and cutaway view. Different acceleration profiles of the oscillating magnet were created by varying the current in the solenoids. **C.** The three oddball conditions consisted of the ‘Opposite pull oddball’ condition (Right pull oddballs after Left pull common and Left pull oddballs after Right pull common), the ‘Pull oddball after neutral’ condition (Left and Right pull oddballs after Neutral common), and the ‘Neutral oddball’ condition (Neutral oddballs after Left and Right common). **D.** Experimental setup showing participant holding the unattached vibrating device in their right hand using a pinch grip whilst high-density EEG was recorded. **E.** Diagram showing relative location of the 129 electrodes. Electrodes from face, ears and neck (shown in grey) were excluded from main analysis due to artefacts.

### Participants

We recruited 15 participants (10 males, 5 females, mean age = 33.33 yrs, SD = 7.33 yrs). All participants were right handed. The sample size was chosen based on previous asymmetric vibration and somatosensory oddball EEG studies (Akatsuka et al., 2007; Spackman et al., 2007; Restuccia et al., 2009; Amemiya and Gomi, 2016). Experiments were undertaken with the understanding and written consent of each participant in accordance with the Code of Ethics of the World Medical Association (Declaration of Helsinki), and with the NTT Communication Science Laboratories Research Ethics Committee approval.

### Procedure

Vibration stimuli were generated by a solenoid actuator within a small device held in a pinch grip. Vibration is generated by a magnet anchored between a pair springs surrounded by a pair of solenoids (Fig. 1B.). The magnet oscillated (58.8Hz) left and right in response to current passing through the solenoids. Leftward acceleration of the solenoid means the fingers receive a rightwards force (and vice versa). By varying the current in the solenoids we generated either symmetric or asymmetric left-right acceleration profiles. Under conditions of asymmetry, one force direction is rendered large and brief, while the other is small and prolonged. Due to nonlinearity in the perceptual system, only the larger of the forces is perceived despite the temporally-integrated forces in each direction being approximately equal (Amemiya et al., 2005; Amemiya and Gomi, 2014). When symmetric vibration is used both forces are equal and cancel each other out. Three forms of vibration were used throughout the experiment; asymmetric left (left pull), asymmetric right (right pull) and symmetric vibration, which is referred to as ‘Neutral’.

First, an accuracy test was administered, in which participants received 100ms bursts of vibration and had to discriminate asymmetric (pulling) from symmetric (neutral) vibrations in a two alternative forced choice task. Responses were given with the left hand via keypad. Left and right pulling stimuli were tested in separate blocks (50 randomized trials per block, 25 per condition; block order counterbalanced). A subset of participants (n = 9) were also required to discriminate between left and right pulling stimuli under the same task conditions.

The main oddball task consisted of vibration stimuli delivered with a randomized ISI of 800ms-1100ms (Fig. 1A.). Vibration stimulus duration was always 100ms. In each block, one vibration pattern (Left, Right or Neutral) was the common stimulus (80% of trials) and the other two were the oddballs (each 10% of trials; total oddball = 20%). Trials were pseudorandomized, such that the first trial of every block was a common stimulus and that every oddball was followed by a common stimulus. Each block consisted of 200 trials (common = 160, oddball A = 20, oddball B= 20). There were 15 blocks in total which were randomized and counterbalanced across participants (5 blocks for each of the 3 block types, defined according to the common stimulus, i.e. Left, Right and Neutral). Thus, in total there were 9 conditions, composed of three common stimuli conditions (Left, Right, Neutral, 800 trials per condition) and 6 oddball stimuli conditions (120 trials per condition). Oddball conditions were grouped into three conditions (Fig. 1C.): ‘Opposite pull oddballs’ (Right oddballs during Left common and Left oddballs during Right common), ‘Pull oddball after neutral’ (Right oddballs during Neutral common and Left oddballs during Neutral common), and ‘Neutral oddball’ (Neutral oddballs during Left common and Neutral oddballs during Right common).

Participants were informed at the start of each block which stimulus was the common and which two were the oddball. They were instructed to pay attention to all stimuli and silently count the number of oddballs. At the end of each block they reported their estimate for the number of oddballs by responding to options presented on screen. Thus, they always simultaneously responded to two oddball conditions, helping to ensure that their effort levels were well controlled across conditions. Participants were naive to the purpose of the experiment when asked directly at the end of testing. The experiment lasted ~2.5 hours.

### Analysis

#### Behavioral data

Accuracy on the pre-test pulling direction discrimination task was determined for each participant by taking the sum of correctly identified pulling and neutral stimuli as a percentage of the total number of trials. Left and Right pulling conditions were calculated separately and compared via paired sample t-test. In the subset of participants (n = 9) who also completed a Left vs. Right pull discrimination block, we calculated the percentage correct in the same manner and compared this value to the mean of the Left vs. Neutral and Right vs. Neutral values via paired sample t-test. Oddball counting error was calculated for each block of the main task by taking the absolute of the estimated number of oddballs minus the actual number of oddballs. Oddball counting error was compared across participants via Wilcoxon signed-rank test. We compared blocks where the left pull was the common stimulus to blocks where right pull was the common stimulus, and we compared the average of these two blocks to blocks in which neutral was the common stimulus.

From the two behavioral tasks we extracted 4 variables that were to be used in covariate analyses with the EEG data: pre-test Pull vs Neutral discrimination (mean of left vs neutral and right vs neutral block), mean oddball counting error (i.e. across all blocks), Left/right common oddball counting error (mean of performance on blocks where left and right pull were the common stimulus), and Neutral common oddball counting error.

#### EEG pre-processing

EEG data were pre-processed using EEGlab (Delorme and Makeig, 2004) and custom Matlab (2017a) scripts. The data were down sampled to 250Hz for storage purposes. We re-referenced the data to the left and right mastoid electrodes and applied a bandpass (FIR 0.1 – 90Hz) and notch filter (48-52Hz). ICA components reflecting blinks, eye movements, heart, large EMG and electrical artefacts were then removed. Epochs were extracted (−200 – 700ms) and baselined (−100 – 0ms) for each participant. Electrodes from the face and side of the head below the ear were removed due to muscle activity artifacts in some participants, leaving a total of 93 electrodes, covering the entire scalp (Fig. 1E.). We removed trials still displaying artefacts via whole brain threshold (+/-80μV) and by applying the ERPlab step function algorithm to frontal electrodes (window size = 200ms, step size = 50ms, threshold = 50μV). The mean percentage of trials rejected per participant was 9.62%. ERPs were averaged across conditions and smoothed using a low pass filter (second order Butterworth, cutoff 30Hz).

#### EEG analysis in surface space using SPM

We analyzed the EEG data in two ways. Firstly, to avoid the bias inherent to picking electrodes and time widows, we used SPM 12 for M/EEG (Litvak et al., 2011) to analyze scalp data across the response window and then to perform source reconstructions of scalp activity. SPM controls for multiple comparisons using Random Field Theory (RFT), which is effective because of the temporal and spatial smoothness of EEG data (Kilner and Friston, 2010). Statistical parametric maps were created for each participant in each condition by interpolating from all electrodes into two-dimensional sensor space across the response window (0-500ms post stimulus onset), thus creating a 3D characterization of the ERP (16mm x 16mm x 0ms smoothing).

To determine if ‘Opposite pull oddball’ produced a larger response than the ‘Pull oddball after neutral’ conditions, it was only necessary to perform a paired t-test (1-tailed) because both oddball conditions used the same common stimulus condition (i.e. common pull). We also ran the same t-contrast using our behavioral variables as covariates. However, we also wanted to check if there were any differences between the ‘Neutral oddball’ condition and the other two oddball conditions, for which we needed to use a partitioned error approach (random effects analysis), combined with two separate 2×2 within-subject’s ANOVAs with factors of oddball condition (Neutral oddball vs Opposite pull oddball or Pull oddball after neutral) and stimulus type (oddball vs common). For these analyses image files (SPM maps; NIFTI) were transformed into four sets of differential effects (overall effect, main effect of condition, main effect of type, condition x type interaction) for each participant (1^st^ level contrasts), which were then entered into four separate one-sample t-tests (2^nd^ level contrasts; for details see Franz et al., 2020). Of these contrasts, only the condition x type interaction was of interest because this contrast showed the effect of the oddball condition, whilst controlling for the common stimulus. For all scalp activity contrasts we used a threshold of p < 0.001 uncorrected and clusters were only included if they met the more stringent p < 0.05 family-wise cluster threshold.

#### EEG source localization

To locate the possible cortical origins of activity detected on the scalp we ran SPM 3-D source reconstruction, using a group inversion approach (COH, 0-500ms, Hanning taper, 0-256Hz) to compensate for head anatomy and sensor noise variation (Litvak and Friston, 2008). An MNI template was used to construct the mesh, coregistration used the nasion and bilateral preauricular points as fiducials, and a forward model was created with the Boundary Elements Model (BEM). NIFTI (source-level) images (8mm smoothing) were extracted using a time window derived from the ‘Opposite pull oddball’ vs ‘Pull oddball after neutral’ scalp analysis (264-320ms). To better refine the location of the activity, NIFTI images were subjected to a paired sample t-test with a general threshold set at P < 0.05 uncorrected, and selected the top cluster of activity (i.e. the cluster that contained the highest peak t-values). Due to the problem of circularity, this statistical test was used purely to better locate the already observed scalp effect (Oh et al., 2020), and to negate the issue of central attraction during source analysis, whereby at the group-level sources can tend to accumulate in biologically implausible central regions of the brain.

To better understand the location of our cluster of pulling related activity, we compared it to the origin of the P50 generated in response to the Neutral common stimuli (an ERP independent of the main ‘Opposite pull oddball’ vs ‘Pull oddball after neutral’ comparison), since the P50 is known to originate in SI (Allison et al., 1992). For this visualization we ran the same SPM group inversion but using a window of −100-100ms and contrasted (paired sample t-test) the baseline period (−100-0ms) with the P50 window (40-64ms) in the Neutral common condition, threshold at P < 0.0006 uncorrected. The threshold was chosen so that the top cluster contained approximately the same number of voxels (2018 voxels) as the ‘pulling related activity’ cluster (2012 voxels).

The location of these clusters of brain activity was compared using the SPM anatomy toolbox (Eickhoff et al., 2005), which provides a list of brain areas ranked according to the likelihood that the observed activity originates within their probabilistically defined boundaries. We considered the top five areas to be representative of the cluster origin, given the spatial limitations of EEG. Ratios (Table 1) are calculated automatically by the toolbox for each area by dividing the mean probability at cluster location by the mean probability across the entire probability map of the brain. Higher values indicate location more towards the center of the area.

**Table 1:** Cluster locations for pull related activity (Opposite pull oddballs vs Pull oddball after neutral) and Neutral common P50 activity, based on maximum probability maps. Estimates of activated areas were based on clusters of 2012 (p < 0.05) and 2018 (p < 0.0006) voxels respectively. Only the five areas calculated to be the most likely origin for activity area shown, sorted according to percent of the cluster volume found in each area. Note that high ratio values indicate higher probabilities that the cluster had an origin in the specific brain area (see methods for details).

| Contrast | Assignment based on Maximum Probability Map | Percent of Cluster volume in Area | Percent of Area activated by Cluster | Ratio |
| --- | --- | --- | --- | --- |
| Pull related activity | Area 7A (SPL) | 11.6 | 13.5 | 1.13 |
|  | Area 2 | 8 | 15.5 | 1.01 |
|  | Area hIP2 (IPS) | 7.2 | 25.7 | 1.77 |
|  | Area hIP6 (IPS) | 5.9 | 15.4 | 1.33 |
|  | Area hIP3 (IPS) | 4.6 | 11 | 0.81 |
| Neutral common P50 | Area 2 | 15.8 | 30.6 | 1.23 |
|  | Area 3b | 11.7 | 16.6 | 1.28 |
|  | Area 7A (SPL) | 9.3 | 10.9 | 1.48 |
|  | Area 4p | 6.6 | 24.9 | 1.55 |
|  | Area 3a | 5.2 | 23.8 | 1.37 |

#### Traditional ERP analysis

We also used a traditional ERP approach using ERPlab (Lopez-Calderon and Luck, 2014), in which we selected electrode locations and time windows based on previous research (Allison et al., 1992; Kekoni et al., 1997; Akatsuka et al., 2005; Shen et al., 2018), wide ERP windows were favored to avoid biasing conditions where the ERPs were flattened due to greater onset variability (Luck, 2014). Epochs were averaged for each of the 9 conditions and oddball difference waves were calculated by subtracting the activity of each stimulus when it was acting as the common stimulus from the activity of the same stimulus when it was acting as an oddball (Pulvermüller et al., 2006). Left and right directional versions of each oddball difference wave were averaged together to give the final experimental condition (Opposite pull oddball), and two other oddball conditions (‘Pull oddball after neutral’ and ‘Neutral oddball’), oddball difference waves.

Mean amplitude of the P50 (30 - 70ms), N140 (100 - 150ms), P200 (150 – 250ms) and P3b (250 – 500ms) event related potentials (ERPs) were quantified from the epoched and difference wave data for each condition. We calculated the onset latency of all ERPs by calculating the point where the signal reached 50% of the peak value within each time window. In line with previous literature, P50 analysis was based on the 6 electrodes surrounding P3, N140 analysis was based on the 5 electrodes surrounding F3, while P200 and P3b analysis was based on the 5 electrodes surrounding Cz. ERP measures were compared across conditions via paired sample t-tests. Our main comparison concerned the ERP responses in the ‘Opposite pull oddball’ condition compared to the ‘Pull oddball after neutral’ condition, however we also compared the ‘Opposite pull oddball’ condition to the ‘Neutral oddball’ condition and compared the ‘Pull oddball after neutral’ condition to the ‘Neutral oddball’ condition.

## Results

### Pulling related activity 280ms post-stimulus over the parietal lobe

We combined a tactile oddball task, in which uncommon target stimuli must be detected from a stream of common stimuli, with asymmetric (left/right pulling) and symmetric vibration (neutral) stimuli (Fig. 1.). ERPs from all conditions can be seen in Figure 2. The purpose of our main contrast, comparing the ‘Opposite pull oddball’ condition and ‘Pull oddball after neutral’ condition (Fig. 1C.) was to find brain activity specific to having a directional pulling expectation violated by a different directional pull (i.e. expect left but get right pull). We analyzed the entire response window for significant clusters in an unbiased manner. This revealed a cluster of significant activity (264-320ms) that peaked over the left parietal cortex (280ms post-stimulus onset) and extended anteriorly to cover part of the left frontal lobe by ~300ms post-stimulus onset (Figs. 5A–5C).

**Figure 2.**
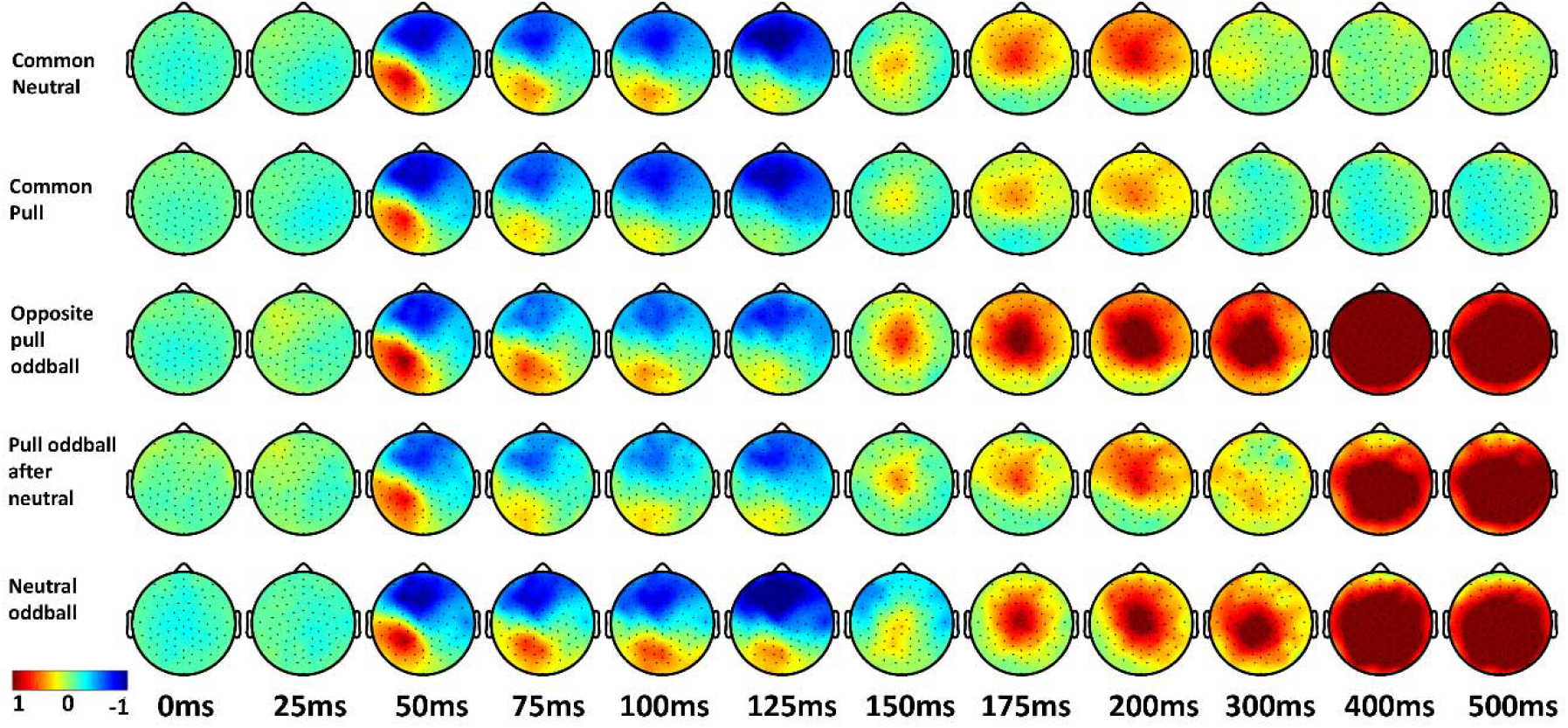
Instantaneous ERP amplitude across conditions. The amplitude (μV) of scalp activity across time in the two common stimulus conditions: ‘Common neutral’ and ‘Common pull’ (mean of Left and Right common), and in the three Oddball conditions: ‘Opposite pull oddball’ (mean of Left oddball after right common and Right oddball after Left common), ‘Pull oddball after neutral’ (mean of Left oddball after Neutral common and Right oddball after Neutral common), and ‘Neutral oddball’ (mean of Neutral oddball after Left common and Neutral oddball after Right common).

### Pulling related activity is spatiotemporally distinct from early processing in sensory and frontal regions

An early sensory-frontal account of the pulling sensation predicts pulling-related amplitude enhancements of the N140. However, the reverse was observed and we did not find evidence that the N140 indexes the pulling sensation. (Fig. 3C–3D; Tables 2 & 3). If the N140 indexed the pulling sensation it should have been present in the ‘Opposite pull oddballs’ condition difference waves (Fig. 3D; red line) and larger than the other two oddball conditions (i.e. more negative). In fact, the mean difference wave from contralateral frontal sites was slightly *positive* in the ‘Opposite pull oddballs’ condition and did not differ from that seen in the ‘Pull oddball after neutral’ condition (0.19μV vs 0.19μV; t (14) = −0.099, p = 0.923, Cohen’s d = −0.02). Indeed, only in the ‘Neutral oddball’ condition was the difference wave negative during the N140 window, with the N140 being significantly larger than that observed for the ‘Opposite pull oddballs’ condition (−0.17μV vs 0.19μV; t (14) = 2.595, p = 0.021, Cohen’s d = 0.84) and the ‘Pull oddball after neutral’ condition (−0.17μV vs 0.19μV; t (14) = 2.599, p = 0.021, Cohen’s d = 0.98). There were no differences in N140 onset latency across conditions (Tables 2 & 3).

**Figure 3.**
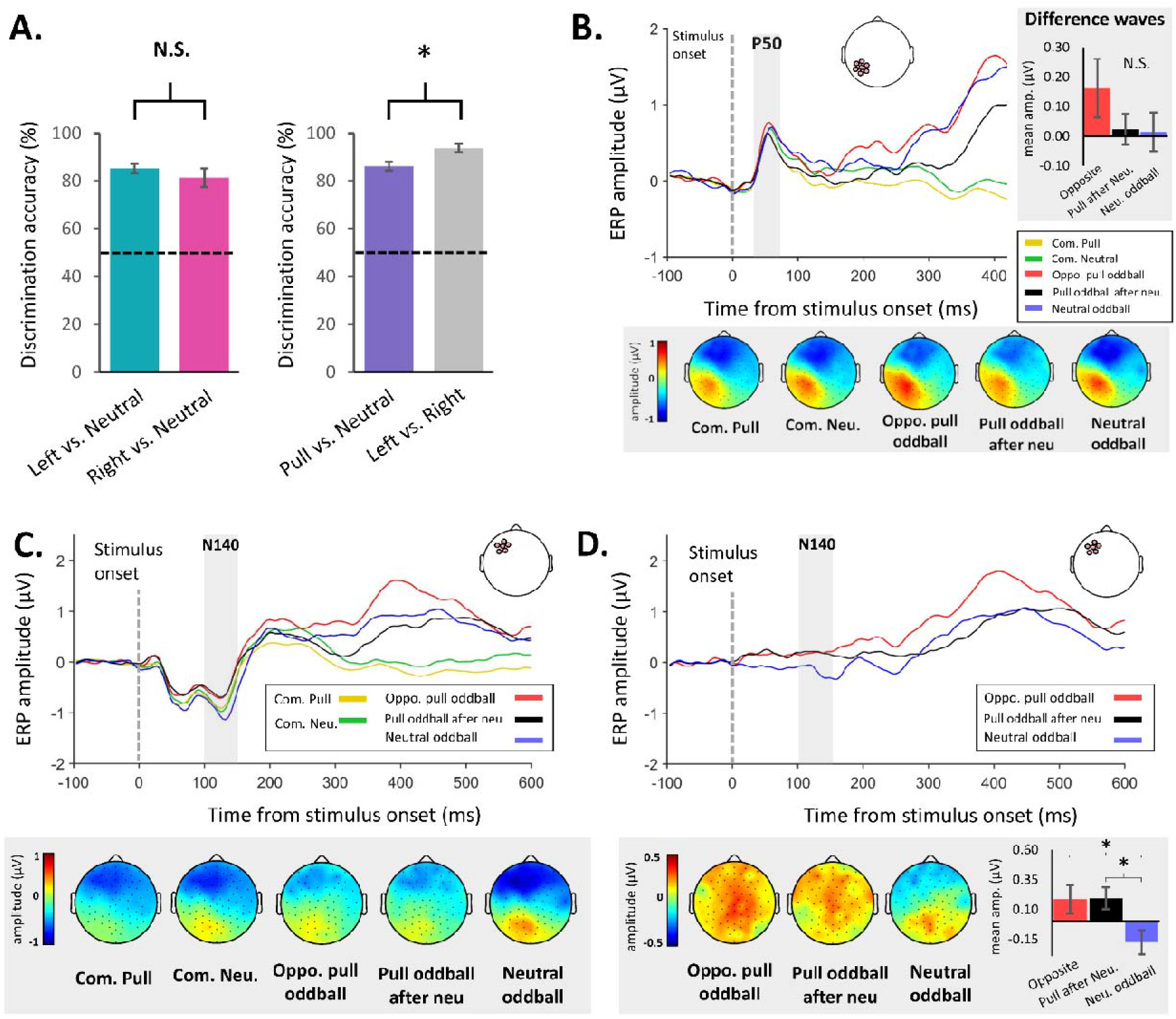
Behavioral, P50 and N140 ERP results. **A.** Group mean pre-test pulling discrimination accuracy was high in all conditions, indicating that all experimental stimuli could clearly be perceived. There was no difference in participants’ ability to discriminate Left and Right pulls (asymmetric vibration) from Neutral stimuli (symmetric vibration), but performance was better in a subset (n = 9) when discriminating Left from Right pulls as opposed from discriminating either of the pulling stimuli from the Neutral stimulus. * *p* < 0.05, N.S. = Not Significant. **B.** Group mean P50 amplitude and scalp maps (30-70ms) for all conditions. When comparing difference waves across the three oddball conditions over the contralateral parietal cortex there was no difference in mean amplitude. N.S. = Not Significant. **C.** Group mean N140 amplitude and scalp maps (100-150ms) for all conditions. **D.** Difference waves from contralateral frontal electrodes showing larger N140 for the ‘Neutral oddball’ condition compared to the ‘Opposite pull oddball’ and Pull oddball after neutral’ condition. * *p* < 0.05, N.S. = Not Significant.

**Table 2:** Onset latency and mean amplitude for P50, N140, P200 and P3b ERPs derived from difference waves for the three oddball conditions. Group (n=15) mean and (SD) values shown. Note that the onset could not always be identified when using difference waves. So for P50 onset latency n = 13 in the Opposite oddball condition and n = 14 in the Neutral oddball condition, for N140 onset latency n = 13 in the Opposite oddball condition and n = 11 in the Pull after Neutral condition, and for P200 onset latency n = 13 in the Pull oddball after neutral condition and n = 14 in the Neutral oddball condition.

| Condition | Latency (ms) |  |  |  | Mean amplitude (µV) |  |  |  |
| --- | --- | --- | --- | --- | --- | --- | --- | --- |
|  | P50 | N140 | P200 | P3b | P50 | N140 | P200 | P3b |
| **Opposite oddball** | 42.68<br>(10.9) | 118.77<br>(12.1) | 173.03<br>(23.88) | 335.42<br>(40.43) | 0.16<br>(0.38) | 0.19<br>(0.46) | 0.68<br>(0.63) | 2.07<br>(1.45) |
| **Pull oddball after neutral** | 38.02<br>(12.2) | 115.49<br>(16.32) | 193.91<br>(25.19) | 382.84<br>(47.87) | 0.02<br>(0.2) | 0.19<br>(0.36) | 0.27<br>(0.5) | 1.25<br>(1.17) |
| **Neutral Oddball** | 45.96<br>(9.96) | 110.41<br>(15.87) | 179.31<br>(19.91) | 338.76<br>(47.16) | 0.01<br>(0.25) | -0.17<br>(0.39) | 0.18<br>(0.63) | 1.7<br>(1.45) |

**Table 3:** Statistical comparison using paired sample t-test of onset latency and mean amplitude for P50, N140, P200 and P3b ERPs comparing the three oddball conditions. Shown are t-values, with p-values in parenthesis (* p < 0.05, ** p < 0.01, *** p < 0.001). DF = 14, except when onset latency could not be identified. P50 latency DF = 12 for Oppo. vs. Pull after neu. and DF = 13 for other two comparisons. N140 latency DF = 8 for Oppo. vs. Pull after neu., DF = 12 for Oppo. vs. Neu. Oddball, and DF = 10 for Pull after neu. vs. Neu. Oddball. P200 latency DF = 12 for Oppo. vs. Pull after neu., DF = 13 for Oppo. vs. Neu. Oddball, DF = 11 for Pull after neu. vs. Neu. Oddball.

| Comparison | Latency |  |  |  | Mean amplitude |  |  |  |
| --- | --- | --- | --- | --- | --- | --- | --- | --- |
|  | P50 | N140 | P200 | P3b | P50 | N140 | P200 | P3b |
| **Oppo. vs. Pull after neu.** | 0.874<br>(0.399) | 0.172<br>(0.867) | -2.388<br>(0.034*) | -3.84<br>(0.002**) | 1.455<br>(0.168) | -0.099<br>(0.923) | 2.818<br>(0.014*) | 2.499<br>(0.026*) |
| **Oppo. vs. Neu. Oddball** | -0.296<br>(0.773) | 1.024<br>(0.326) | -0.799<br>(0.439) | -0.412<br>(0.686) | 1.164<br>(0.264) | 2.595<br>(0.021*) | 2.240<br>(0.042*) | 1.104<br>(0.288) |
| **Pull after neu. vs. Neu. Oddball** | -1.496<br>(0.158) | 0.531<br>(0.607) | 1.684<br>(0.120) | 5.221<br>(<0.001***) | 0.143<br>(0.889) | 2.599<br>(0.021*) | 0.479<br>(0.639) | -2.118<br>(0.053) |

We did not observe any differences in the P50 amplitude or latency across oddball conditions (Fig. 3B.; Tables 2 & 3). An early, sensory-frontal account predicts higher feedforward activity in SI, which we did not observe. Nevertheless, it is difficult to conclude anything from this null result, which could simply be due to noise in the data.

Next, we sought to determine whether pulling related activity was spatially distinct from early sensory activity. The results must be interpreted cautiously owing to the inverse problem and poor spatial acuity of the EEG signal (Grech et al., 2008). Group inversion of the pulling-related scalp activity (Fig. 5D, red and green patches; Table 1) suggested an origin in the left parietal cortex, corresponding to the postcentral sulcus, superior parietal lobule (SPL) and the intraparietal sulcus (IPS). The SPM anatomy toolkit indicated that the cluster was centered on the IPS (Fig. 5.; Table 1).

For comparison we analyzed the approximate location of SI using the Neutral common condition P50 activity, since the P50 has been shown to have its main origin inside SI (Allison et al., 1992). This cluster was found to be located somewhat anterior, though partially overlapping with, our pulling related cluster of activity (Fig. 5D, blue and green patches; Table 1). The SPM anatomy toolkit indicated there was some P50 activity in the postcentral sulcus and SPL, as with the pulling related activity. However, unlike the pulling related activity, the P50 cluster was not strongly represented in the IPS, and instead was represented in the postcentral gyrus, consistent with the approximate location of the SI hand area (Holmes et al., 2019).

Taken together the results argue against an early sensory-frontal account of the pulling sensation. The N140 was, contrary to the sensory-frontal account, attenuated for pulling oddballs and enhanced for neutral oddballs. Pulling related activity occurred later (280ms; *see also P200 and P3b results below*) and was centered on the parietal association cortex, overlapping with, but spatially distinct from the site of early sensory activity.

### Lack of significant activity when comparing to neutral oddballs

We did not observe any significant clusters of activity when the ‘Neutral oddball’ condition was compared to either of the other two oddball conditions using the SPM scalp analysis. This may in part have been due to the large P3b generated by Neutral oddballs (Fig. 4.) which obscured any differences between the ‘Neutral oddball’ and ‘Opposite pull oddball’ conditions (*see below and discussion*).

**Figure 4.**
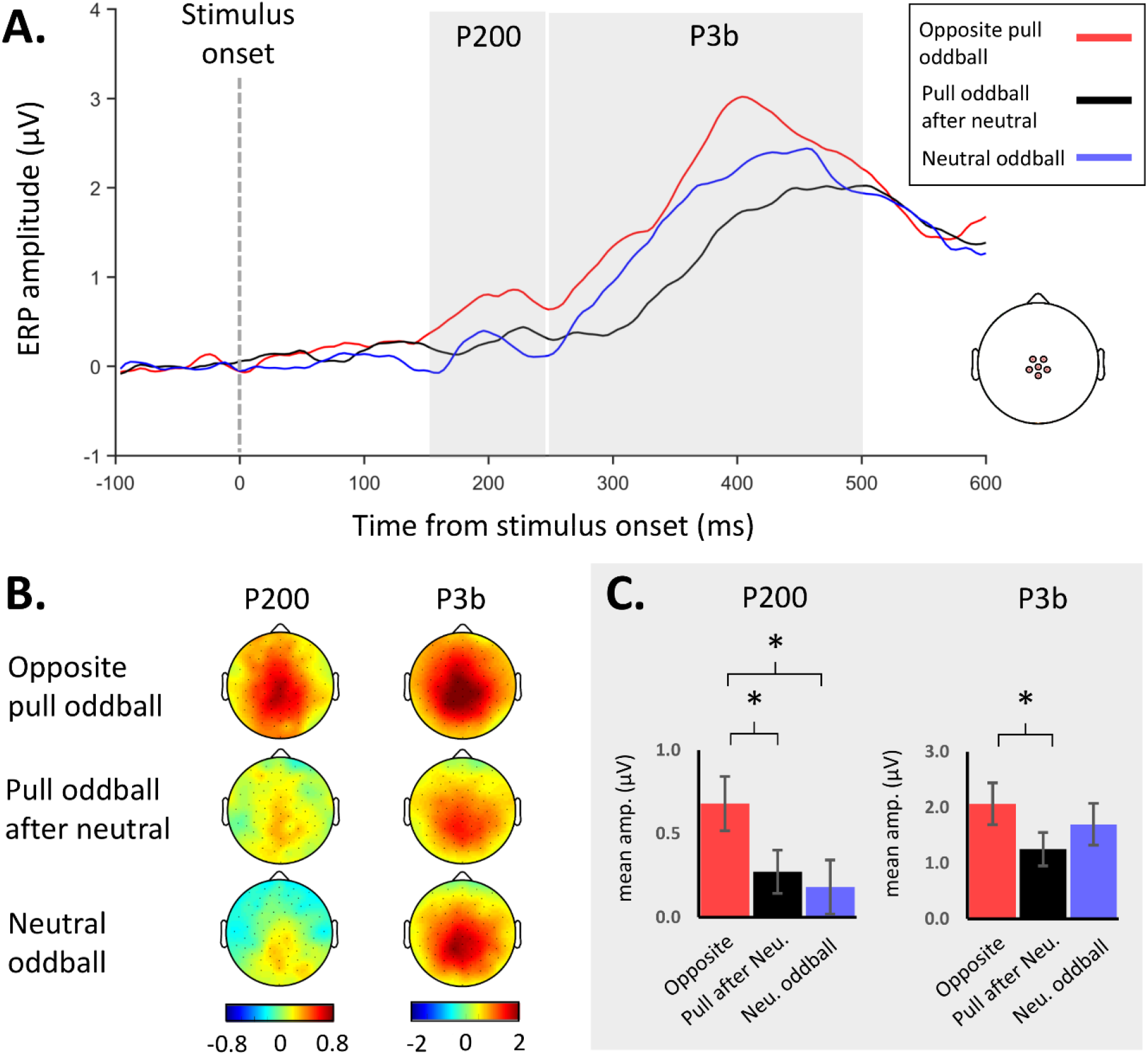
P200 and P3b oddball effects across conditions. **A.** Group average ERP oddball difference waves (subtraction of activity related to common stimuli) from central electrodes shown for the three oddball conditions. P200 (150-250ms) and P3b (250-500ms) windows shown in grey. **B.** Group average difference wave scalp activity for the P200 and P3b in each of the three oddball conditions. **C.** Group average P200 amplitude was significantly larger in the ‘Opposite pull oddball’ condition than both the other two oddball conditions, while P3b activity was larger in the ‘Opposite pull oddball’ condition than the ‘Pull oddball after neutral’ condition. * *p* < 0.05.

**Figure 5.**
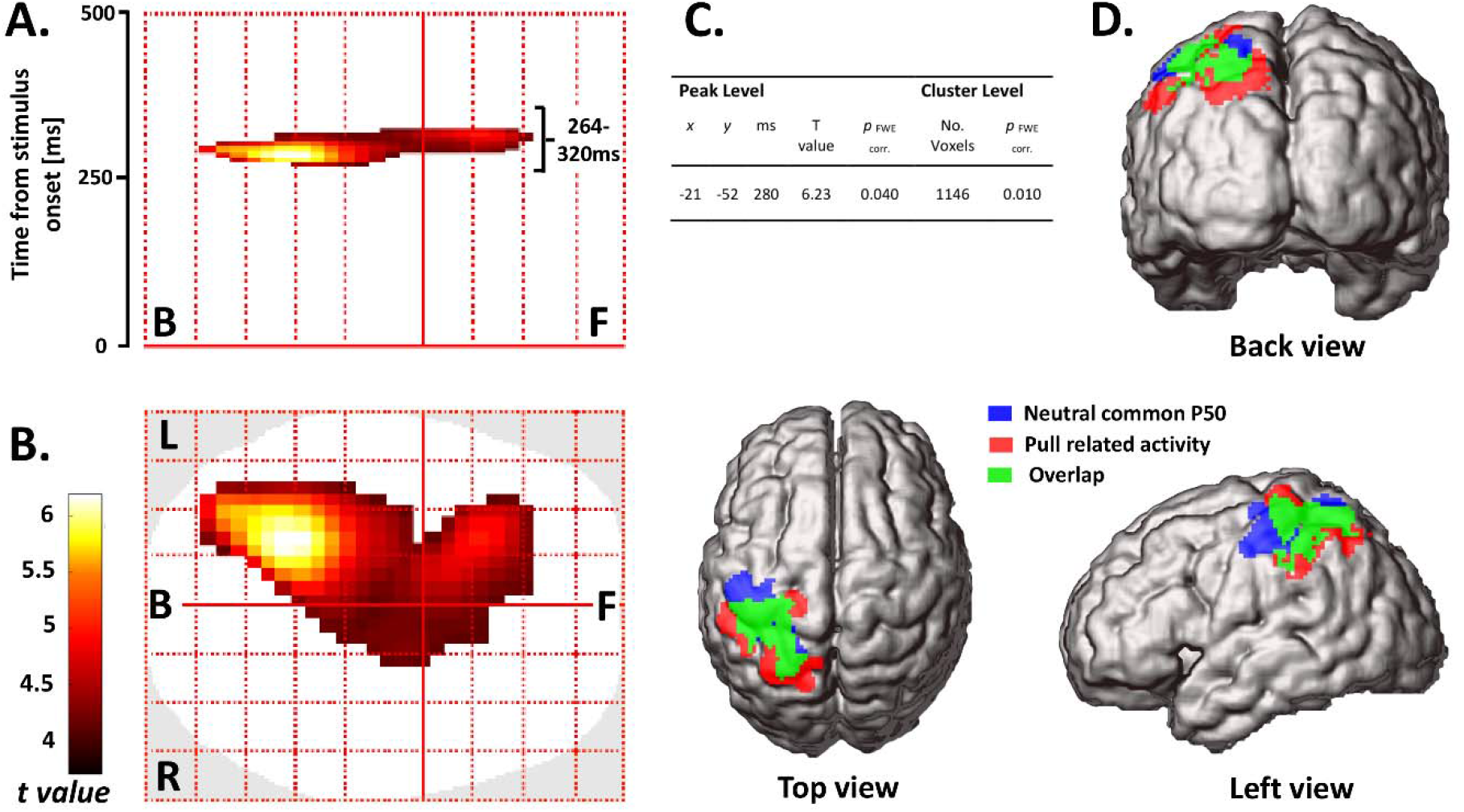
Brain activity associated with the pulling sensation. **A-C**. Results of SPM topographical analysis in sensor space when contrasting the ‘Opposite pull oddball’ and ‘Pull oddball after neutral’ conditions. A cluster of significant activity (p < 0.001 uncorrected, p < 0.05 FWE cluster threshold) was observed (264-320ms after stimulus onset), peaking at 280ms over the left parietal cortex and extending anteriorly. In table, *x* position is positive-going left to right, *y* position is positive-going from posterior to anterior. **D.** Group inversion (p < 0.05 uncorrected) suggested the significant scalp activity originated from the left parietal cortex. To clarify the location of this activity we conducted a separate group inversion using the Neutral common condition, in the P50 time window (40-64ms). This was overlaid on the previously identified pulling related activity, and showed a more anterior distribution, close to the SI hand area.

### Pulling related activity associated with earlier and larger P200 and P3b responses

The ‘Opposite pull oddball’ condition was associated with larger amplitude P200 responses than the ‘Pull oddball after neutral’ condition (0.68μV vs 0.27μV; t (14) = 2.818, p = 0.014, Cohen’s d = 0.72; Fig 4; Tables 2 & 3) and the ‘Neutral oddball’ condition (0.68μV vs 0.18μV; t (14) = 2.24, p = 0.042, Cohen’s d = 0.79). P200 latency was also shorter for the ‘Opposite pull oddball’ condition compared to the ‘Pull oddball after neutral’ condition (173.03ms vs 193.91ms; t (12) = −2.388, p = 0.034, Cohen’s d = −0.85), but not when the Opposite pull oddball’ condition was compared to the ‘Neutral oddball condition’ (t (13) = −0.799, p = 0.439).

For the P3b the ‘Opposite pull oddball’ condition was also associated with larger amplitude (2.07μV vs 1.25μV; t (14) = 2.499, p = 0.026, Cohen’s d = 0.62; Fig. 4; Tables 2 & 3) and shorter latency (335.42ms vs 382.84ms; t(14) = −3.84, p = 0.002, Cohen’s d = − 1.07) responses compared to the ‘Pull oddball after neutral’ condition. No significant differences in P3b amplitude or latency were observed when comparing the ‘Opposite pull oddball’ condition with the ‘Neutral oddball’ condition (Fig. 4; Tables 2 & 3.). This was likely because the P3b was larger than expected in the ‘Neutral oddball’ condition. Indeed, for the P3b, compared to the ‘Pull after neutral oddball’ condition, the ‘Neutral oddball’ condition was associated with earlier ERP onset latencies (t(14) = 5.221, p < 0.001, Cohen’s d = 0.93) and a trend towards larger mean amplitudes (t(14) = −2.118, p = 0.053, Cohen’s d = 0.34; Fig. 4; Tables 2 & 3.).

### Opposite direction pulls easier to discriminate behaviorally

We tested participants’ ability to discriminate between the three different types of stimuli (Left/Right pulls and Neutral). All participants were able to clearly feel the pulling sensation (Fig. 3A). A subset of tested participants was better at discriminating Left vs. Right than discriminating Left/Right vs Neutral (93.78% (SD = 7.03) vs. 86.22% (SD = 6.83); t(8) = −2.591, p = 0.032, Cohen’s d = 1.09; Fig. 3A. right panel). This finding was expected, given that Left/Right discrimination is similar to the ‘Opposite pull oddball’ condition, which was associated with the strongest brain response across conditions. We did not find any difference in pre-test pulling discrimination accuracy when comparing Left vs. Neutral to Right vs. Neutral (85.33% (SD = 7.62) vs. 81.45% (SD = 14.9); t(14) = − 1.439, p = 0.172; Fig. 3A. left panel). When asked, participants reported that neutral stimuli (symmetric vibration) were subjectively similar to asymmetric vibration, aside from the absence of the pulling sensation.

Responses were not biased to the left or right. Participants did not differ in their propensity to respond left as opposed to neutral (47.6% vs. 52.4%; t(14) = 1.103, p = 0.288) or right as opposed to neutral (53.07% vs. 46.93%; t(14) = −1.667, p = 0.118) during the pre-test discrimination task. Likewise, there was no difference when left and right were compared directly, either using the discrimination from neutral conditions (t(14) = 0.544, p = 0.595) or when left and right pulling sensations were discriminated from one another directly (52.22% vs 47.78%; t(8) = −0.989, p = 0.352).

During the main task, oddball counting error did not differ when comparing Left Common to Right Common blocks (5.49 (SD = 3.23) vs 4.73 (SD = 2.51); Z = −0.483, p = 0.631) or when comparing mean Left/Right common blocks to Neutral common blocks (5.11 (SD = 2.34) vs 5.73 (SD = 2.34); Z = 1.079, p =0.28). This lack of difference suggests there were no major differences in performance level, effort or attention across block types, which, if present, could have confounded our interpretation of the EEG results.

The behavioral measures were added as covariates in our SPM analyses, but the results did not substantially change (Table 4), ruling out the possibility that our clusters of significant activity were artefacts of extremes of task performance.

**Table 4:** Results of the SPM sensor space contrast comparing the ‘Opposite pull oddball condition’ and ‘Pull oddball after neutral’ condition, using behavioral measures as covariates. Threshold was set at p < 0.001 uncorrected and only clusters that passed family-wise (FWE) cluster threshold of p < 0.1 were included, *x* position is positive-going left to right, *y* position is positive-going from posterior to anterior.

| Behavioral covariate | Peak Level | | | T value | $p_{\text{FWE corr.}}$ | Cluster Level | |
| --- | --- | --- | --- | --- | --- | --- | --- |
| | x | y | ms | | | No. Voxels | $p_{\text{FWE corr.}}$ |
| Pull vs Neutral discrimination | -21 | -52 | 280 | 6.59 | 0.036 | 757 | 0.030 |
| Mean oddball count error | -21 | -52 | 280 | 6.01 | 0.067 | 537 | 0.061 |
| Left/Right common, oddball count error | -21 | -52 | 280 | 6.10 | 0.062 | 568 | 0.055 |
| Neutral common, oddball count error | -21 | -46 | 280 | 6.41 | 0.043 | 897 | 0.020 |

## Discussion

Research using monkeys (Salimi et al., 1999; Backlund Wasling et al., 2008; Fortier-Poisson et al., 2016) and work addressing peripheral tactile processing in humans (Birznieks et al., 2001; Pruszynski and Johansson, 2014) suggested that the rapid extraction of force vectors is possible. Drawing on this, and work detailing the SI, SII and PFC circuitry involved in tactile frequency discrimination (Romo and Salinas, 2003), we formulated a sensory-frontal account of the pulling sensation. This account holds that spatial processing is not necessary for pulling sensations, and that directional pulls can be identified with reference to stored patterns of activity. In contrast, a spatial account argues that such processing is not sufficient, and that activity in the somatosensory association cortex is instead required for the pulling sensation to emerge. We used high-density EEG, combined with an illusory pulling sensation embedded within an oddball task, and found evidence that supported this latter, somatosensory association cortex account.

### Evidence against an early sensory-frontal account of pulling sensations

The sensory-frontal account predicted that pulling sensations should enhance the N140, either directly, because of shared underlying mechanisms, or indirectly because of the redirection of attention. Direct enhancement was suggested by the fact that the N140 generators overlap with the SII and PFC circuitry involved in tactile frequency discrimination (Allison et al., 1992; Frot et al., 1999; Valeriani et al., 2001; de Lafuente and Romo, 2006), and because the N140 indexes somatosensory awareness (Auksztulewicz et al., 2012; Auksztulewicz and Blankenburg, 2013; Forschack et al., 2020; Schröder et al., 2021) and texture perception (Genna et al., 2018), processes closely related to the pulling sensation. Indirectly, the N140 is implicated in the detection of oddballs (Kekoni et al., 1997; Andrew et al., 2020), and with both endogenous and exogenous tactile attention (Nakajima and Imamura, 2000). As such, if the pulling sensation emerged upstream of N140 generators, greater attention should have been directed towards the stimulus, resulting in downstream N140 enhancement (Forster and Eimer, 2004).

However, we did not find any such pulling-related enhancements of the N140. We observed the opposite: pulling sensations slightly attenuated the N140, while neutral oddball stimuli actually produced a larger N140 than pulling oddballs. This result suggests that the sensory-frontal account is unlikely to be correct and that pulling sensations emerged via alternate mechanisms. Conversely, neutral oddballs (symmetric vibration), which participants reported being subjectively similar to asymmetric vibration aside from the absence of the pulling sensation, were apparently processed in a manner more similar to how vibration frequency is discriminated.

Our N140 findings constrain theoretical accounts, since they suggest that the pulling sensation does not emerge during the initial stages of somatosensory processing. Consistent with this, we did not observe any pulling-related P50 effects, although little can be concluded from this null result. Indeed, our results do not contradict earlier research involving monkeys that reported rapid SI processing of tangential forces (Salimi et al., 1999). Early processing in SI is undoubtedly necessary, but as our results show, likely not sufficient for the pulling sensation to emerge. Moreover, we found that later pulling-related activity (264-320ms) did overlap with the posterior portion of SI. SI may contribute to awareness of the pulling sensation via reentrant activity from the parietal lobe (Auksztulewicz et al., 2012; Meador et al., 2017).

### A somatosensory association cortex account of the pulling sensation

Pulling related activity occurred 264-320ms post-stimulus, beginning and peaking (280ms) over the contralateral parietal lobe. Source localization indicated an origin in the postcentral sulcus, SPL, and IPS, posterior to an independent localization of SI. This pattern of activity is consistent with a spatial account of the pulling sensation, specifically that feeling a directional tactile pull depends on the integration of body location processing with processing of the force vector orientation and direction in space.

Unlike other bodily illusions involving illusory external forces (De Havas et al., 2017, 2018), asymmetric vibration produces a clear sensation that the hand is being pulled yet is stationary. The brain therefore needs to determine that the hand is not moving despite an apparent external force. The postcentral sulcus may contribute to the pulling sensation by providing input regarding hand position, since this is a key somatosensory area for proprioception (Soechting and Flanders, 1989; Cohen and Andersen, 2002; London and Miller, 2013; Chowdhury et al., 2020) and is involved in generating vibration-based proprioceptive body illusions (Ehrsson et al., 2005).

Our parietal cluster of pulling-related activity was centered on the IPS, which may contribute to the pulling sensation by extracting the orientation of the illusory force vector, as with the orientation of graspable objects (Hadjikhani and Roland, 1998; Frey et al., 2005; Van Boven et al., 2005; Wacker et al., 2011; Leoné et al., 2015). IPS activity might also be related to anticipatory grip control (Ehrsson et al., 2003; van Polanen et al., 2020), indicating that such control can be decoupled from the actual need to adjust grip strength.

The SPL is involved in spatial cognition (Colby and Goldberg, 1999; Sack, 2009), body location processing (Graziano et al., 2000; Felician et al., 2004) and transformations into body centered reference frames (Lacquaniti et al., 1995; Gallivan et al., 2009). To determine where a pull is directed, and to dissociate pulls from merely moving tactile stimuli (Lin and Kajola, 2003; Oh et al., 2017), processes in the SPL could map an extracted force vector in hand or externally-centered coordinates.

The poor spatial acuity of EEG means we cannot be certain which of these brain areas represent the true loci of pulling related activity. Nevertheless, we can be confident about the temporal evolution of the pulling sensation, which emerged 280ms post-stimulus and modified the P200. Pulling-related activity can therefore be temporally dissociated from the predominantly feedforward processing in SI and SII (< 70ms), as well as from the initial engagement of the parietal cortex (70-100ms), which, during oddball tasks, is likely related to identifying the unusualness of the stimulus (Huang et al., 2005). Instead, the timing of the pulling related activity is similar to other somatosensory illusions, such as the rubber hand illusion, which has recently been found to be associated with parietal and frontal activity 200-300ms post-stimulus (Rao and Kayser, 2017; Guterstam et al., 2019). Of more direct relevance, the temporal characteristics of the pulling sensation are consistent with tactile remapping. Tactile remapping, whereby a force vector is transformed into bodily centered coordinate system, depends on activity in the SPL and IPS (Azañón et al., 2010; Ritterband-Rosenbaum et al., 2014; Heed et al., 2015), takes place after initial somatosensory processing has been completed, and is closely associated with the P200 (Longo et al., 2012; Bufalari et al., 2014). Thus, converging lines of evidence point towards a spatial, parietal cortex account of the pulling sensation.

### Processing related to the violation and updating of sensory expectations

Comparing opposite direction oddballs to the same oddballs after neutral stimuli was designed to isolate activity specific to having directional pulling expectations violated by directional pulling sensations. We are therefore acknowledging an inherently predictive account of perception (Berthoz, 2000; Friston, 2005). However, it is difficult to exclude activity related to the process of comparison itself (Garrido et al., 2009; Camalier et al., 2019).

Two caveats must be addressed. Firstly, within a Bayesian framework, our main result, that opposite pull oddballs produced larger parietal activity than the same oddballs in the context of neutral common stimuli, could be argued to be due to neutral common stimuli forming weaker priors than pulling stimuli, which in turn would produce a smaller response when the priors were confounded by the oddball. This argument, however, is not wholly convincing because opposite pull oddballs produced a larger P200 response than neutral oddballs, despite having matched common stimuli, and thus the same priors. Additionally, neutral common stimuli produced slightly larger ERPs than pulling common stimuli, suggesting comparable or greater salience, and rendering weaker priors doubtful.

A second, related caveat, is that the parietal lobe activity we observed might reflect the contradiction of an expected pulling direction by *any* tactile stimulus, since we did not observe the same pattern of activity when contrasting opposite pull oddballs with neutral oddballs. This result was likely due to the inherent uncertainty of the neutral stimulus when acting as an oddball, a factor known to amplify long-latency ERPs (Stern et al., 2010; Furl and Averbeck, 2011; Kopp et al., 2016), and explaining the large P3b found in the Neutral oddball condition.

After the pulling sensation has been generated and its direction determined, stimulus classification and memory updating processes can begin; processes indexed by the P3b (Polich, 2007). Earlier and larger P3b responses for opposite direction oddballs were probably observed because the extracted oddball pulling direction was maximally different from the common stimulus extracted direction (Miltner et al., 1989; Nakajima and Imamura, 2000).

### Conclusion

Our findings suggest the sensation of being pulled emerges through parietal lobe activity 280ms post-stimulus, related to processing proprioception, tactile orientation and peripersonal space. This first step towards a spatiotemporally precise account of the pulling sensation will aid the development of handheld vibration devices for gaming, navigation and guiding the visually impaired (Amemiya and Sugiyama, 2010; Takamuku et al., 2016; Gomi et al., 2019) and may shed light on the role of pulling sensations in parent-infant communication and developmental disorders characterized by somatosensory deficits (Cascio, 2010). Understanding the neural mechanisms of the pulling sensation delineates its commonalities and differences with other sensorimotor processes, such as tactile motion detection, grasping control and bodily awareness, furthering a more complete account of parietal lobe function.

## Acknowledgements

This work was supported by Grants-in-Aid for Scientific Research (JP16H06566) from Japan Society for the Promotion of Science to HG, and by a research contract between NTT and UCL.

## Author contributions

JDH, SI and HG conceived the study and designed the experiments. JDH and SI collected the data. JDH, SB and HG conceived the data analysis. JDH analyzed the data. JDH, SB and HG wrote the manuscript. All authors provided comments and approved the manuscript.

## References

Akatsuka K, Wasaka T, Nakata H, Inui K, Hoshiyama M, Kakigi R (2005) Mismatch responses related to temporal discrimination of somatosensory stimulation. Clin Neurophysiol Off J Int Fed Clin Neurophysiol 116:1930–1937.

Akatsuka K, Wasaka T, Nakata H, Kida T, Kakigi R (2007) The effect of stimulus probability on the somatosensory mismatch field. Exp Brain Res 181:607–614.

Allison T, McCarthy G, Wood CC (1992) The relationship between human long-latency somatosensory evoked potentials recorded from the cortical surface and from the scalp. Electroencephalogr Clin Neurophysiol 84:301–314.

Amemiya T, Ando H, Maeda T (2005) Virtual force display: direction guidance using asymmetric acceleration via periodic translational motion. In: First Joint Eurohaptics Conference and Symposium on Haptic Interfaces for Virtual Environment and Teleoperator Systems. World Haptics Conference, pp 619–622.

Amemiya T, Gomi H (2014) Distinct Pseudo-Attraction Force Sensation by a Thumb-Sized Vibrator that Oscillates Asymmetrically. In: Haptics: Neuroscience, Devices, Modeling, and Applications (Auvray M, Duriez C, eds), pp 88–95 Lecture Notes in Computer Science. Berlin, Heidelberg: Springer.

Amemiya T, Gomi H (2016) Active Manual Movement Improves Directional Perception of Illusory Force. IEEE Trans Haptics 9:465–473.

Amemiya T, Maeda T (2008) Asymmetric Oscillation Distorts the Perceived Heaviness of Handheld Objects. IEEE Trans Haptics 1:9–18.

Amemiya T, Sugiyama H (2010) Orienting Kinesthetically: A Haptic Handheld Wayfinder for People with Visual Impairments. ACM Trans Access Comput 3:6:1-6:23.

Andrew D, Ibey RJ, Staines WR (2020) Transient inhibition of the cerebellum impairs change-detection processes: Cerebellar contributions to sensorimotor integration. Behav Brain Res 378:112273.

Auksztulewicz R, Blankenburg F (2013) Subjective rating of weak tactile stimuli is parametrically encoded in event-related potentials. J Neurosci Off J Soc Neurosci 33:11878–11887.

Auksztulewicz R, Spitzer B, Blankenburg F (2012) Recurrent Neural Processing and Somatosensory Awareness. J Neurosci 32:799–805.

Azañón E, Longo MR, Soto-Faraco S, Haggard P (2010) The Posterior Parietal Cortex Remaps Touch into External Space. Curr Biol 20:1304–1309.

Backlund Wasling H, Lundblad L, Löken L, Wessberg J, Wiklund K, Norrsell U, Olausson H (2008) Cortical processing of lateral skin stretch stimulation in humans. Exp Brain Res 190:117–124.

Badde S, Heed T (2016) Towards explaining spatial touch perception: Weighted integration of multiple location codes. Cogn Neuropsychol 33:26–47.

Berthoz A (2000) The Brain’s Sense of Movement. Harvard University Press.

Birznieks I, Jenmalm P, Goodwin AW, Johansson RS (2001) Encoding of direction of fingertip forces by human tactile afferents. J Neurosci Off J Soc Neurosci 21:8222–8237.

Brainard DH (1997) The Psychophysics Toolbox. Spat Vis 10:433–436.

Bufalari I, Di Russo F, Aglioti SM (2014) Illusory and veridical mapping of tactile objects in the primary somatosensory and posterior parietal cortex. Cereb Cortex N Y N 1991 24:1867–1878.

Camalier CR, Scarim K, Mishkin M, Averbeck BB (2019) A Comparison of Auditory Oddball Responses in Dorsolateral Prefrontal Cortex, Basolateral Amygdala, and Auditory Cortex of Macaque. J Cogn Neurosci 31:1054–1064.

Cascio CJ (2010) Somatosensory processing in neurodevelopmental disorders. J Neurodev Disord 2:62–69.

Chowdhury RH, Glaser JI, Miller LE (2020) Area 2 of primary somatosensory cortex encodes kinematics of the whole arm. eLife 9.

Cohen YE, Andersen RA (2002) A common reference frame for movement plans in the posterior parietal cortex. Nat Rev Neurosci 3:553–562.

Colby CL, Goldberg ME (1999) Space and attention in parietal cortex. Annu Rev Neurosci 22:319–349.

De Havas J, Gomi H, Haggard P (2017) Experimental investigations of control principles of involuntary movement: a comprehensive review of the Kohnstamm phenomenon. Exp Brain Res.

De Havas J, Ito S, Haggard P, Gomi H (2018) Low Gain Servo Control During the Kohnstamm Phenomenon Reveals Dissociation Between Low-Level Control Mechanisms for Involuntary vs. Voluntary Arm Movements. Front Behav Neurosci 12 Available at: https://www.frontiersin.org/articles/10.3389/fnbeh.2018.00113/full [Accessed May 30, 2018].

de Lafuente V, Romo R (2006) Neural correlate of subjective sensory experience gradually builds up across cortical areas. Proc Natl Acad Sci U S A 103:14266–14271.

Delorme A, Makeig S (2004) EEGLAB: an open source toolbox for analysis of single-trial EEG dynamics including independent component analysis. J Neurosci Methods 134:9–21.

Desmedt JE, Tomberg C (1989) Mapping early somatosensory evoked potentials in selective attention: critical evaluation of control conditions used for titrating by difference the cognitive P30, P40, P100 and N140. Electroencephalogr Clin Neurophysiol 74:321–346.

Driver J, Spence C (1998) Cross-modal links in spatial attention. Philos Trans R Soc B Biol Sci 353:1319–1331.

Ehrsson HH, Fagergren A, Johansson RS, Forssberg H (2003) Evidence for the Involvement of the Posterior Parietal Cortex in Coordination of Fingertip Forces for Grasp Stability in Manipulation. J Neurophysiol 90:2978–2986.

Ehrsson HH, Kito T, Sadato N, Passingham RE, Naito E (2005) Neural Substrate of Body Size: Illusory Feeling of Shrinking of the Waist. PLOS Biol 3:e412.

Eickhoff SB, Stephan KE, Mohlberg H, Grefkes C, Fink GR, Amunts K, Zilles K (2005) A new SPM toolbox for combining probabilistic cytoarchitectonic maps and functional imaging data. NeuroImage 25:1325–1335.

Felician O, Romaiguère P, Anton J-L, Nazarian B, Roth M, Poncet M, Roll J-P (2004) The role of human left superior parietal lobule in body part localization. Ann Neurol 55:749–751.

Forschack N, Nierhaus T, Müller MM, Villringer A (2020) Dissociable neural correlates of stimulation intensity and detection in somatosensation. NeuroImage 217:116908.

Forster B, Eimer M (2004) The attentional selection of spatial and non-spatial attributes in touch: ERP evidence for parallel and independent processes. Biol Psychol 66:1–20.

Fortier-Poisson P, Langlais J-S, Smith AM (2016) Correlation of fingertip shear force direction with somatosensory cortical activity in monkey. J Neurophysiol 115:100–111.

Fortier-Poisson P, Smith AM (2016) Neuronal activity in somatosensory cortex related to tactile exploration. J Neurophysiol 115:112–126.

Franz M, Schmidt B, Hecht H, Naumann E, Miltner WHR (2020) Suggested deafness during hypnosis and simulation of hypnosis compared to a distraction and control condition: A study on subjective experience and cortical brain responses. PloS One 15:e0240832.

Frey SH, Vinton D, Norlund R, Grafton ST (2005) Cortical topography of human anterior intraparietal cortex active during visually guided grasping. Brain Res Cogn Brain Res 23:397–405.

Friston K (2005) A theory of cortical responses. Philos Trans R Soc Lond B Biol Sci 360:815–836.

Frot M, Rambaud L, Guénot M, Mauguière F (1999) Intracortical recordings of early pain-related CO2-laser evoked potentials in the human second somatosensory (SII) area. Clin Neurophysiol Off J Int Fed Clin Neurophysiol 110:133–145.

Furl N, Averbeck BB (2011) Parietal cortex and insula relate to evidence seeking relevant to reward-related decisions. J Neurosci Off J Soc Neurosci 31:17572–17582.

Gallivan JP, Cavina-Pratesi C, Culham JC (2009) Is that within reach? fMRI reveals that the human superior parieto-occipital cortex encodes objects reachable by the hand. J Neurosci Off J Soc Neurosci 29:4381–4391.

Garrido MI, Kilner JM, Stephan KE, Friston KJ (2009) The mismatch negativity: a review of underlying mechanisms. Clin Neurophysiol Off J Int Fed Clin Neurophysiol 120:453–463.

Genna C, Oddo C, Fanciullacci C, Chisari C, Micera S, Artoni F (2018) Bilateral cortical representation of tactile roughness. Brain Res 1699:79–88.

Gomi H, Ito S, Tanase R (2019) Innovative mobile force display: Buru-Navi. In: International Display Workshops. Sapporo, Japan.

Graziano MS, Cooke DF, Taylor CS (2000) Coding the location of the arm by sight. Science 290:1782–1786.

Grech R, Cassar T, Muscat J, Camilleri KP, Fabri SG, Zervakis M, Xanthopoulos P, Sakkalis V, Vanrumste B (2008) Review on solving the inverse problem in EEG source analysis. J NeuroEngineering Rehabil 5:25.

Guterstam A, Collins KL, Cronin JA, Zeberg H, Darvas F, Weaver KE, Ojemann JG, Ehrsson HH (2019) Direct Electrophysiological Correlates of Body Ownership in Human Cerebral Cortex. Cereb Cortex N Y N 1991 29:1328–1341.

Hadjikhani N, Roland PE (1998) Cross-modal transfer of information between the tactile and the visual representations in the human brain: A positron emission tomographic study. J Neurosci Off J Soc Neurosci 18:1072–1084.

Heed T, Buchholz VN, Engel AK, Röder B (2015) Tactile remapping: from coordinate transformation to integration in sensorimotor processing. Trends Cogn Sci 19:251–258.

Hernández A, Nácher V, Luna R, Zainos A, Lemus L, Alvarez M, Vázquez Y, Camarillo L, Romo R (2010) Decoding a perceptual decision process across cortex. Neuron 66:300–314.

Holmes NP, Tamè L, Beeching P, Medford M, Rakova M, Stuart A, Zeni S (2019) Locating primary somatosensory cortex in human brain stimulation studies: experimental evidence. J Neurophysiol 121:336–344.

Huang M-X, Lee RR, Miller GA, Thoma RJ, Hanlon FM, Paulson KM, Martin K, Harrington DL, Weisend MP, Edgar JC, Canive JM (2005) A parietal-frontal network studied by somatosensory oddball MEG responses, and its cross-modal consistency. NeuroImage 28:99–114.

Johansson RS, Flanagan JR (2009) Coding and use of tactile signals from the fingertips in object manipulation tasks. Nat Rev Neurosci 10:345–359.

Johansson RS, Hger C, Bäckström L (1992a) Somatosensory control of precision grip during unpredictable pulling loads. III. Impairments during digital anesthesia. Exp Brain Res 89:204–213.

Johansson RS, Riso R, Häger C, Bäckström L (1992b) Somatosensory control of precision grip during unpredictable pulling loads. I. Changes in load force amplitude. Exp Brain Res 89:181–191.

Kekoni J, Hämäläinen H, Saarinen M, Gröhn J, Reinikainen K, Lehtokoski A, Näätänen R (1997) Rate effect and mismatch responses in the somatosensory system: ERP-recordings in humans. Biol Psychol 46:125–142.

Kida T, Nishihira Y, Hatta A, Wasaka T (2003) Somatosensory N250 and P300 during discrimination tasks. Int J Psychophysiol Off J Int Organ Psychophysiol 48:275–283.

Kilner JM, Friston KJ (2010) TOPOLOGICAL INFERENCE FOR EEG AND MEG. Ann Appl Stat 4:1272–1290.

Kopp B, Seer C, Lange F, Kluytmans A, Kolossa A, Fingscheidt T, Hoijtink H (2016) P300 amplitude variations, prior probabilities, and likelihoods: A Bayesian ERP study. Cogn Affect Behav Neurosci 16:911–928.

Lacquaniti F, Guigon E, Bianchi L, Ferraina S, Caminiti R (1995) Representing spatial information for limb movement: role of area 5 in the monkey. Cereb Cortex N Y N 1991 5:391–409.

Leoné FTM, Monaco S, Henriques DYP, Toni I, Medendorp WP (2015) Flexible Reference Frames for Grasp Planning in Human Parietofrontal Cortex. eNeuro 2.

Lin Y-Y, Kajola M (2003) Neuromagnetic somatosensory responses to natural moving tactile stimulation. Can J Neurol Sci J Can Sci Neurol 30:31–35.

Litvak V, Friston K (2008) Electromagnetic source reconstruction for group studies. NeuroImage 42:1490–1498.

Litvak V, Mattout J, Kiebel S, Phillips C, Henson R, Kilner J, Barnes G, Oostenveld R, Daunizeau J, Flandin G, Penny W, Friston K (2011) EEG and MEG data analysis in SPM8. Comput Intell Neurosci 2011:852961.

London BM, Miller LE (2013) Responses of somatosensory area 2 neurons to actively and passively generated limb movements. J Neurophysiol 109:1505–1513.

Longo MR, Musil JJ, Haggard P (2012) Visuo-tactile integration in personal space. J Cogn Neurosci 24:543–552.

Lopez-Calderon J, Luck SJ (2014) ERPLAB: an open-source toolbox for the analysis of event-related potentials. Front Hum Neurosci 8:213.

Luck SJ (2014) An Introduction to the Event-Related Potential Technique. MIT Press.

Meador KJ, Revill KP, Epstein CM, Sathian K, Loring DW, Rorden C (2017) Neuroimaging somatosensory perception and masking. Neuropsychologia 94:44–51.

Miltner W, Johnson R, Braun C, Larbig W (1989) Somatosensory event-related potentials to painful and non-painful stimuli: effects of attention. Pain 38:303–312.

Nakajima Y, Imamura N (2000) Relationships between attention effects and intensity effects on the cognitive N140 and P300 components of somatosensory ERPs. Clin Neurophysiol Off J Int Fed Clin Neurophysiol 111:1711–1718.

Oh H, Custead R, Wang Y, Barlow S (2017) Neural encoding of saltatory pneumotactile velocity in human glabrous hand. PLOS ONE 12:e0183532.

Oh Y, Chesebrough C, Erickson B, Zhang F, Kounios J (2020) An insight-related neural reward signal. NeuroImage 214:116757.

Panarese A, Edin BB (2011) Human ability to discriminate direction of three-dimensional force stimuli applied to the finger pad. J Neurophysiol 105:541–547.

Polich J (2007) Updating P300: an integrative theory of P3a and P3b. Clin Neurophysiol Off J Int Fed Clin Neurophysiol 118:2128–2148.

Pruszynski JA, Flanagan JR, Johansson RS (2018) Fast and accurate edge orientation processing during object manipulation. eLife 7.

Pruszynski JA, Johansson RS (2014) Edge-orientation processing in first-order tactile neurons. Nat Neurosci 17:1404–1409.

Pulvermüller F, Shtyrov Y, Ilmoniemi RJ, Marslen-Wilson WD (2006) Tracking speech comprehension in space and time. NeuroImage 31:1297–1305.

Rao IS, Kayser C (2017) Neurophysiological Correlates of the Rubber Hand Illusion in Late Evoked and Alpha/Beta Band Activity. Front Hum Neurosci 11:377.

Restuccia D, Zanini S, Cazzagon M, Del Piero I, Martucci L, Della Marca G (2009) Somatosensory mismatch negativity in healthy children. Dev Med Child Neurol 51:991–998.

Ritterband-Rosenbaum A, Hermosillo R, Kroliczak G, van Donkelaar P (2014) Hand position-dependent modulation of errors in vibrotactile temporal order judgments: the effects of transcranial magnetic stimulation to the human posterior parietal cortex. Exp Brain Res 232:1689–1698.

Romo R, Salinas E (2003) Flutter Discrimination: neural codes, perception, memory and decision making. Nat Rev Neurosci 4:203–218.

Sack AT (2009) Parietal cortex and spatial cognition. Behav Brain Res 202:153–161.

Salimi I, Brochier T, Smith AM (1999) Neuronal activity in somatosensory cortex of monkeys using a precision grip. III. Responses to altered friction perturbations. J Neurophysiol 81:845–857.

Schröder P, Nierhaus T, Blankenburg F (2021) Dissociating Perceptual Awareness and Postperceptual Processing: The P300 Is Not a Reliable Marker of Somatosensory Target Detection. J Neurosci 41:4686–4696.

Shen G, Smyk NJ, Meltzoff AN, Marshall PJ (2018) Using somatosensory mismatch responses as a window into somatotopic processing of tactile stimulation. Psychophysiology 55:e13030.

Shinozaki N, Yabe H, Sutoh T, Hiruma T, Kaneko S (1998) Somatosensory automatic responses to deviant stimuli. Brain Res Cogn Brain Res 7:165–171.

Soechting JF, Flanders M (1989) Errors in pointing are due to approximations in sensorimotor transformations. J Neurophysiol 62:595–608.

Spackman LA, Boyd SG, Towell A (2007) Effects of stimulus frequency and duration on somatosensory discrimination responses. Exp Brain Res 177:21–30.

Stern ER, Gonzalez R, Welsh RC, Taylor SF (2010) Updating beliefs for a decision: neural correlates of uncertainty and underconfidence. J Neurosci Off J Soc Neurosci 30:8032–8041.

Takamuku S, Amemiya T, Ito S, Gomi H (2016) Design of Illusory Force Sensation for Virtual Fishing. Trans Hum Interface Soc 18:87–94.

Tanabe T, Yano H, Iwata H (2018) Evaluation of the Perceptual Characteristics of a Force Induced by Asymmetric Vibrations. IEEE Trans Haptics 11:220–231.

Tappeiner HW, Klatzky RL, Unger B, Hollis R (2009) Good vibrations: Asymmetric vibrations for directional haptic cues. In: World Haptics 2009 - Third Joint EuroHaptics conference and Symposium on Haptic Interfaces for Virtual Environment and Teleoperator Systems, pp 285–289.

Valeriani M, Fraioli L, Ranghi F, Giaquinto S (2001) Dipolar source modeling of the P300 event-related potential after somatosensory stimulation. Muscle Nerve 24:1677–1686.

Van Boven RW, Ingeholm JE, Beauchamp MS, Bikle PC, Ungerleider LG (2005) Tactile form and location processing in the human brain. Proc Natl Acad Sci U S A 102:12601–12605.

van Polanen V, Rens G, Davare M (2020) The role of the anterior intraparietal sulcus and the lateral occipital cortex in fingertip force scaling and weight perception during object lifting. J Neurophysiol 124:557–573.

Wacker E, Spitzer B, Lützkendorf R, Bernarding J, Blankenburg F (2011) Tactile motion and pattern processing assessed with high-field FMRI. PloS One 6:e24860.

